# Microglia are not necessary for maintenance of blood-brain barrier properties in health, but PLX5622 alters brain endothelial cholesterol metabolism

**DOI:** 10.1101/2024.02.20.580754

**Authors:** CP Profaci, SS Harvey, K Bajc, TZ Zhang, AZ Zhang, KM Nemec, GL McKinsey, A Longworth, TP McMullen, TD Arnold, DA Lawson, FC Bennett, D Davalos, R Daneman

**Affiliations:** Department of Pharmacology, University of California, San Diego. La Jolla, CA. USA; Department of Neurosciences, University of California, San Diego. La Jolla, CA. USA; Department of Psychiatry, Perelman School of Medicine, University of Pennsylvania, Philadelphia, PA, USA; Department of Pediatrics, University of California, San Francisco. San Francisco, CA. USA; Department of Physiology and Biophysics, University of California, Irvine. Irvine, CA. USA; Department of Neurosciences, Cleveland Clinic Lerner Research Institute. Cleveland, OH. USA

**Keywords:** blood-brain barrier, microglia, cholesterol, LDLR, CSF1R, PLX5622

## Abstract

Microglia are resident immune cells of the central nervous system, yet their functions far exceed those related to immunology. From pruning neural synapses during development to preventing excessive neural activity throughout life, microglia are intimately involved in the brain’s most basic processes. Studies have reported a close interaction between microglia and endothelial cells, as well as both helpful and harmful roles for microglia at the blood-brain barrier (BBB) in the context of disease. However, much less work has been done to understand microglia-endothelial cell interactions in the healthy brain. Here, we aim to determine the role of microglia at the healthy BBB. We used the colony-stimulating factor 1 receptor (CSF1R) inhibitor PLX5622 to deplete microglia and analyzed BBB ultrastructure, permeability, and transcriptome. Interestingly, we found that, despite their direct contact with endothelial cells, microglia are not necessary for maintenance of BBB structure, function, or gene expression in the healthy brain. However, we found that PLX5622 treatment alters brain endothelial cholesterol metabolism, and this effect was independent from microglial depletion, suggesting PLX5622 has off-target effects on brain vasculature.

## Introduction

While perivascular, meningeal, and choroid plexus macrophages occupy different niches within the central nervous system (CNS), microglia are the only CNS parenchymal macrophages^1^. Microglia are incredibly dynamic cells, constantly moving their ramified processes to survey their environment^2,3^. They adopt different roles in a context-dependent manner, and their many functions include, but are not limited to, releasing pro- or anti-inflammatory cytokines in response to pathological stimuli^1,4^, phagocytosing dysfunctional synapses^5,6^, sculpting neuronal circuits and myelination in development^5,7–9^, and modulating neuronal activity levels in adulthood^10,11^.

Microglia arise from yolk sac progenitors, traveling to the brain at approximately embryonic day 9.5 (E9.5) in the developing mouse^12^. From the early days of their residency in the brain, microglia are intimately involved in refining neural circuits via synaptic pruning^5,7,8^. Their interaction with CNS vasculature also begins early in development. Juxtavascular microglia migrate along blood vessels as they colonize the developing brain^13^, and microglia interact with endothelial tip cells to promote vascular anastomosis^14^. In the adult brain, approximately 20-30% of microglia make somatic contacts with vasculature^13,15^. Interestingly, this interaction is even higher in the first week of postnatal development in mice and at gestational weeks 18-24 in humans^13^, although the functional reason for this peak is still poorly understood. In addition to somatic contacts with vasculature, microglial processes also make contacts with vasculature, reaching between astrocyte endfeet to touch the endothelial basement membrane^16^.

Despite the physical interaction of microglia and endothelial cells in the healthy CNS, research on this interaction has primarily focused on how microglia impact the blood-brain barrier (BBB) in disease states. Several groups have shown that microglia can actively help repair the BBB after injury. Fernández-López et al. found that, after neonatal stroke, microglial depletion increased hemorrhage in the injured region^17^. Lou et al. demonstrated that, upon two-photon laser injury of a single capillary, microglia immediately extend processes towards injury and help to reseal the BBB^18^. In zebrafish, Liu et al. captured videos of microglia adhering to two ends of an injured vessel, using mechanical forces to pull the endothelial cells back together^19^. However, there is also evidence that microglia accelerate BBB dysfunction in some situations, likely in more extreme cases of neuroinflammation. For instance, microglia form perivascular clusters in EAE, a mouse model of MS. At the peak of EAE, these clustered microglia produce increased levels of reactive oxygen species^20^, known mediators of BBB disruption. It is evident that microglia can play varied roles at the BBB in the context of disease, even having opposite effects within the time course of a single disease. For example, microglia were shown to be initially protective but eventually harmful to BBB integrity during systemic inflammation^21^.

Despite demonstrated functional roles for microglia at the BBB in pathological states, as well as reports that microglia contribute to neurovascular coupling^15,22^, there is no clear understanding of whether microglia regulate the individual properties of the healthy BBB (tight junctions, limited transcytosis, efflux, nutrient transport, limited leukocyte adhesion molecules). In this study, we used a colony-stimulation factor 1 receptor (CSF1R) inhibitor, PLX5622, to deplete microglia and then assessed BBB ultrastructure, permeability, and transcriptome. Manipulation of CSF1 signaling has been a widely used method for probing microglial function, as microglia require constant CSF1 or IL-34 signaling through the CSF1 receptor for survival^23,24^. Pharmacological inhibition of CSF1R or genetic deletion of *Csf1r* or its enhancer is sufficient to deplete microglia quickly and extensively *in vivo*^23,25,26^. While Elmore et al. reported no BBB breakdown upon microglial depletion^25^, they assessed the BBB by imaging the intact brain to detect parenchymal extravasation of Evans Blue dye, a method that would likely only reveal severe BBB disruption. Therefore, it remains unknown whether microglia are required for maintaining more complex aspects of BBB ultrastructure, function, and/or gene expression. Delaney et al. suggest that reduced CSF1R signaling can cause remodeling of BBB tight junctions *in vitro*, and they show evidence of vascular pathology in post-mortem samples from people with a CSF1R mutation^27^. However, a direct link between CSF1R signaling and BBB dysfunction has yet to be clearly demonstrated *in vivo*.

After one month of pharmacological microglial depletion, we assessed BBB ultrastructure via transmission electron microscopy (TEM). To probe barrier permeability, we used two fluorescent tracers with different properties, sodium fluorescein (polar) and rhodamine123 (non-polar). Finally, we acutely purified endothelial cells from control and microglia-deficient mice and performed RNA sequencing to determine the effects of microglial depletion on the endothelial transcriptome. We found that microglia are not required for maintenance of BBB ultrastructure or function, or the expression of genes associated with BBB properties, including tight junctions, transcytosis, efflux transport, solute transport, or leukocyte adhesion. However, to our surprise, we found that PLX5622 treatment increases brain endothelial expression of cholesterol synthesis enzymes and the cholesterol uptake receptor, while decreasing expression of the cholesterol efflux transporter. This phenotype was specific to central nervous system (CNS) endothelial cells and occurred throughout the vascular tree. Finally, through careful time course studies and use of genetic microglial depletion models, we show this effect to be independent of microglial depletion. Together our results show that PLX5622 treatment, but not microglial depletion, increases brain endothelial expression of cholesterol synthesis and uptake machinery.

## Results

### Microglial depletion does not affect ultrastructure of the BBB

To determine whether microglial depletion affects BBB ultrastructure, adult wildtype mice were fed control chow or chow containing 1200 mg/kg of CSF1R inhibitor PLX5622 for one month. We chose a one-month time point to avoid the potential confounding factor of widespread cell death in the early days of drug administration. After one month of diet, mice fed PLX5622 chow exhibited 95% microglial depletion (**Fig 1A-B**). Transmission electron microscopy was used to image blood vessel cross-sections in cortical tissue. Images were analyzed for BBB structural abnormalities, including altered tight junction length and levels of vesicle transport (**Fig 1C-E**). Tight junctions from control mice and microglia-depleted mice were an average length of 385.0 ± 49.38 nm and 480.9 ± 93.58 nm, respectively, a non-significant difference (p=0.400) (**Fig 1D**). Control and microglia-depleted mice had 0.3 ± 0.15 and 0.3 ± 0.12 actively invaginating vesicles per vessel cross-section, respectively (p>0.999) (**Fig 1E**). Taken together, these data suggest that microglia are not required for maintenance of tight junction length or vesicle trafficking at the healthy BBB.

**Figure 1.**
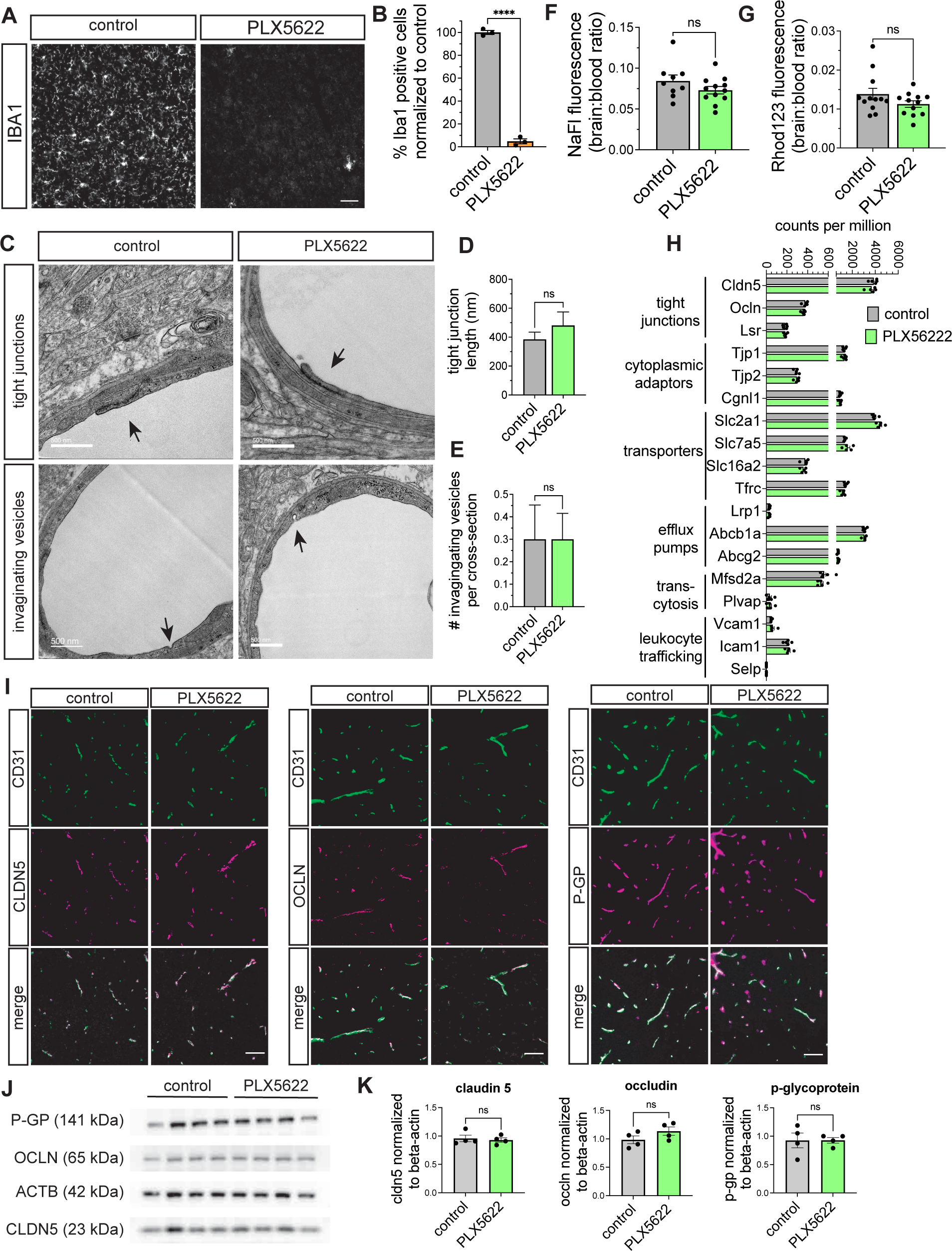
Microglia are not required for BBB properties. 8- to 12-week-old wildtype C57BL/6 mice were put on control or PLX5622 (1200mg/kg) diet for one month. (A) Representative images of 10μm sections stained with anti-IBA1 demonstrate that PLX5622 diet significantly depletes microglia. Scale bar represents 50 μm. (B) Quantification of microglial cell bodies in the cortex. Each data point represents one mouse. Data are represented as a percentage of microglia normalized to the average of control. Error bars represent SEM. n=3 mice per group; p<0.0001, unpaired two-tailed t-test. (C) Representative transmission electron microscopy images of vessel cross-sections in cortical tissue. Images show examples of tight junctions (arrows, top row) and invaginating vesicles (arrows, bottom row). Scale bar represents 500 nm. (D) Quantification of tight junction length in EM images. n=3 mice per group, 20 vessel cross-sections per mouse. Average tight junction length was 480.9 ± 93.58 nm, p=0.400, Mann-Whitney test. Error bars represent SEM. (E) Quantification of invaginating vesicles per vessel cross-section in EM images. n=3 mice per group, 20 vessel cross-sections per mouse. Average number of invaginating vesicles per cross-section was 0.3 ± 0.15 and 0.3 ± 0.12, p>0.999, Mann-Whitney test. Error bars represent SEM. (F) Quantification of permeability to sodium fluorescein. Permeability was calculated as a ratio of [brain]:[blood] fluorescence. There were no group differences in permeability to sodium fluorescein. n=9-12; p=0.183, unpaired two-tailed t-test. Error bars represent SEM. (G) Quantification of permeability to rhodamine 123. Permeability was calculated as a ratio of [brain]:[blood] fluorescence. There were no group differences in permeability to rhodamine 123. n=12; p=0.150, unpaired two-tailed t-test. Error bars represent SEM. (H) Expression of blood-brain barrier genes in acutely isolated endothelial cells. Expression is presented in counts per million; error bars represent SEM. There were no significant differences among any genes associated with classic BBB properties (n=5 mice per group, p-adj > 0.05 for all genes). (I) Representative images of cortex from 10 μm sections stained for CD31 and either claudin5, occludin, or p-glycoprotein (P-GP). There was no observable difference in tight junction or P-GP expression or localization. Scale bar represents 50 μm. (J) Western blot analysis of homogenized cortex for key BBB genes. Membrane was cut and stained for P-GP, occludin, claudin 5, or beta-actin as a loading control. All images are from the same membrane. (K) Quantification of band size. Claudin 5, occludin, and P-GP bands were normalized to the respective beta-actin band in that lane. An unpaired two-tailed t-test was used to test significance. There were no differences in claudin 5 (p=0.709) occludin (p=0.178), or p-glycoprotein (p=0.925). n=4 mice per group. Error bars represent SEM.

### Microglial depletion does not alter BBB permeability

To determine whether microglial depletion alters BBB permeability, adult wildtype mice were fed control or PLX5622 diet for one month. Mice were then injected with either sodium fluorescein or rhodamine 123 to detect BBB permeability (**Fig 1F-G**). Sodium fluorescein is a small, hydrophilic molecule, and its detection in the brain parenchyma is commonly used to measure passive permeability, which might signify disrupted tight junctions between neighboring endothelial cells or increased non-specific transcytosis across endothelial cells. Rhodamine 123, in contrast, is a hydrophobic P-GP/BCRP substrate, and its increased presence in brain parenchyma would be evidence of dysfunctional efflux transport at the BBB. Microglial depletion did not significantly alter the [brain]:[blood] permeability ratio of either sodium fluorescein or rhodamine 123. Control and PLX5622-treated mice exhibited permeability ratios of 0.084 ± 0.007 and 0.073 ± 0.005, respectively (p=0.183) in the sodium fluorescein assay (**Fig 1F**). In the rhodamine 123 assay, control and PLX5622-treated mice exhibited permeability ratios of 0.014 ± 0.001 and 0.011 ± 0.001, respectively (p=0.150) (**Fig 1G**). These data suggest that microglia are not required to maintain the BBB’s restriction of small hydrophilic molecules or hydrophobic efflux substrates.

### Microglial depletion does not alter expression of BBB genes

To explore whether microglial depletion affects the expression of BBB-related genes, we performed bulk RNA sequencing on acutely isolated endothelial cells from mice fed control or PLX5622 chow for one month. We compared mRNA expression levels of genes coding for tight junction proteins, efflux transporters, nutrient transporters, regulators of transcytosis, and leukocyte adhesion molecules. There were no significant differences in the expression of this module of BBB genes (**Fig 1H**). These data were validated by immunohistochemistry and Western blot of key tight junction proteins and efflux transporter (**Fig 1I-K**). There was no significant difference in protein concentration of claudin 5, occludin, or p-glycoprotein between the brains of mice fed control or PLX5622 diet (p=0.709; p=0.178; p=0.986) (**Fig 1K**). In sum, these results suggest that microglia are not necessary for maintaining endothelial expression of genes that modulate key BBB properties in the healthy brain.

### PLX5622 treatment causes upregulation of brain endothelial cholesterol genes

To our surprise, however, we found that PLX5622 treatment increased the expression of a cassette of genes associated with cholesterol metabolism (**Fig 2A**). Specifically, PLX5622 treatment increased brain endothelial expression of a series of 16 cholesterol synthesis enzymes: *Hmgcr*, *Hmgcs1*, *Mvd*, *Mvk*, *Idi1*, *Fdps*, *Fdft1*, *Sqle*, *Lss*, *Cyp51*, *Msmo1*, *Nsdhl*, *Hsd17b7*, *Dhcr7*, *Sc5d*, and *Dhcr24*. PLX5622 also upregulated the cholesterol synthesis regulators *Insig1* and *Srebf2*, as well as the cholesterol uptake receptor *Ldlr*. Conversely, the main cholesterol efflux receptor, *Abca1*, was downregulated in PLX5622-treated mice (**Fig 2A**). These data suggest that PLX5622 treatment leads brain endothelial cells to synthesize and take up cholesterol while reducing cholesterol efflux. To validate this phenotype, we quantified vascular length positive for LDLR in control and PLX5622-treated mice (**Fig 2B-C**). We found that the percentage of LDLR+ vascular length increased from 14.63% ± 1.66 to 73.4% ± 5.56 (**Fig 2C**). All of our sequencing experiments and immunohistochemical quantification were performed with C57BL/6 mice. To rule out strain specificity, we tested the effect of PLX5622 on CBA/J mice and found that, after one week of PLX5622 diet, there was a similarly stark increase in vascular LDLR, increasing from 17.53% ± 4.88 to 61.22% ± 4.26 (**Fig 2D**). Additionally, we validated sequencing results by performing qPCR on vessel fractions from the brains of mice fed control or PLX5622 diet for one month. We found that PLX5622 treatment indeed caused increased expression of *Ldlr* and *Dchr24*, the final enzyme in the cholesterol synthesis pathway (p=0.0033; p=0.0009) (**Fig 2E-F**). These data confirm that, across strains, PLX5622 treatment alters brain endothelial LDLR expression.

**Figure 2.**
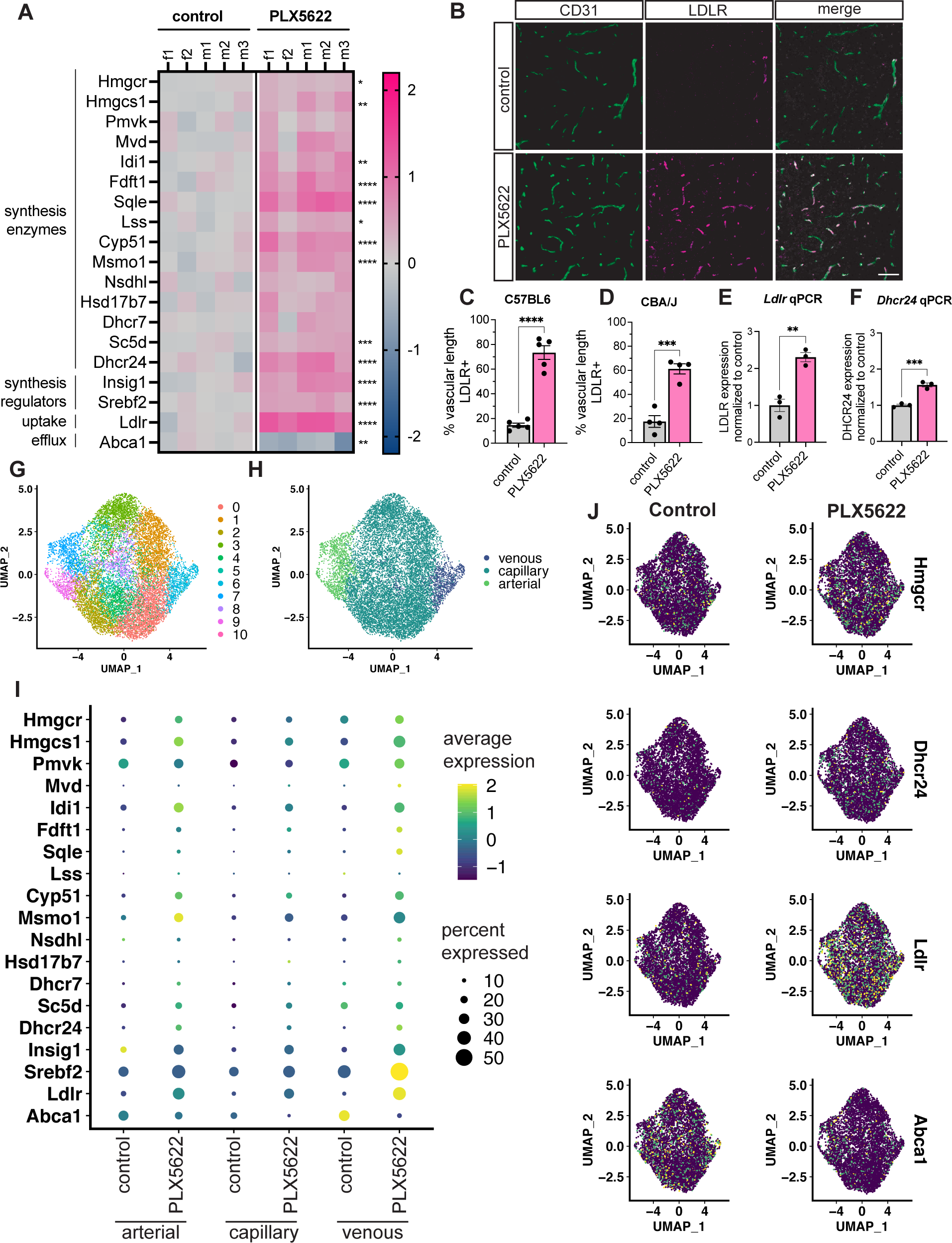
PLX5622 treatment increases brain endothelial expression of cholesterol-related genes. (A) Cholesterol-related gene expression of brain endothelial cells from C57BL/6 mice fed control or PLX5622 (1200 mg/kg) diet for one month. Gene expression for each sample is presented as the log_2_(FC) from the average of control expression, with pink representing an increase and blue representing a decrease. Each column is a sample from one male (m) or female (f) mouse. PLX5622 causes endothelial upregulation of genes coding for cholesterol synthesis enzymes and uptake receptor, and downregulation of the gene coding for the primary cholesterol efflux transporter. Asterisks indicate FDR adjusted p-value (*p-adj<0.05; **p-adj<0.01; ***p-adj<0.001; ****p-adj<0.0001) (B) Representative images of 10μm cortex sections after one month control or PLX5622 diet in adult C57BL/6 mice. Sections were stained with antibodies against CD31 (green, endothelial cells) and LDLR (magenta). PLX5622 diet causes a visible increase in vascular LDLR expression. Scale bar represents 50 μm. (C) Quantification of data shown in (B) examining percentage of CD31+ vascular length that is also LDLR positive. PLX5622 increases percent vascular length that is LDLR+ (n=5; p<0.0001, unpaired two-tailed t-test). Error bars represent SEM. (D) Quantification of vascular LDLR in cortex of 8-to12-week-old CBA/J mice. Mice were put on control or PLX5622 diet for one week. 10μm images of cortex were stained with antibodies against CD31 (green, endothelial cells) and LDLR (magenta) and percentage of CD31+ vascular length that is also LDLR+ was quantified. PLX5622 increases percent vascular length that is LDLR+. n=4; p=0.0005, unpaired two-tailed t-test. Error bars represent SEM. (E) qPCR validation of PLX5622-induced expression changes in *Ldlr*. qPCR was performed on brain vessel fractions. Each data point represents expression levels in vessels isolated from 3 mice. PLX5622 increases *Ldlr* expression 2.3 fold. n=3 samples, p=0.0033, unpaired t-test. Error bars represent SEM. (F) qPCR validation of PLX5622-induced expression changes in *Dhcr24,* the final enzyme in the cholesterol synthesis pathway. qPCR was performed on brain vessel fractions. Each data point represents expression levels in vessels isolated from 3 mice. PLX5622 increased *Dhcr24* expression 1.6 fold. n=3 samples, p=0.0009, unpaired t-test. Error bars represent SEM. (G) UMAP plot showing 11 endothelial cell clusters from single-cell sequencing of brain endothelial cells isolated from adult wildtype mice fed control or PLX5622 diet. For each diet condition, cells were obtained from three brains across two samples. (H) The UMAP plot shown in (G) pseudo-colored based on brain endothelial cell identity as venous (light green), capillary (turquoise), or arterial (dark blue). Identity was determined based on gene expression profiles. Position on the vascular tree was the driving factor of heterogeneity. (I) Dot-plot of brain endothelial cell expression of cholesterol-related genes in control and PLX5622 diet conditions. For each gene, the size of the dot represents the percentage of endothelial cells expressing that gene, and the color of the dot represents average expression on a log scale, with yellow signifying highest expression and purple signifying lowest. Across the arterial-venous axis, the expression of cholesterol synthesis enzymes, synthesis regulators, and uptake receptor were increased. The expression of the cholesterol efflux transporter was decreased. (J) UMAP plots in which brain endothelial cells are colored based on their expression of Hmgcr, Dhcr24, Ldlr, and Abca1. Lighter colors represent higher expression. PLX5622 treatment increased brain endothelial expression of *Hmgcr*, *Dhcr24*, and *Ldlr*, and decreased expression of *Abca1*.

To test whether this effect was occurring throughout the vascular tree, we again put adult mice on control or PLX5622 diet for one month and performed single-cell sequencing on acutely isolated endothelial cells. Eleven clusters of endothelial cells were identified by principal component analysis (**Fig 2G**). The central differences in expression were attributable to the cells’ location along the vascular tree (**Fig 2H**). Analysis of the cholesterol gene cassette revealed the same pattern across arterial, capillary, and venous endothelial cells: an overall increase both in the percent of endothelial cells with detectable expression of cholesterol genes, and in expression level per cell (**Fig 2I-J**). As in our bulk sequencing, the efflux transporter *Abca1* showed the opposite trend as synthesis and uptake genes (**Fig 2I-J**). Together, these data confirm our bulk sequencing findings and demonstrate that PLX5622 alters cholesterol metabolism in brain endothelial cells throughout the vascular tree.

### PLX5622-mediated modulation of cholesterol metabolism in the brain is limited to endothelial cells

The brain is thought to make its cholesterol *de novo*, but brain endothelial cells are not major producers of cholesterol in the brain; this role falls largely to astrocytes and oligodendrocytes. Microglia are also not among the major cholesterol producers and depend on astrocyte-produced cholesterol for survival^24^. We thought perhaps the absence of microglia might change astrocyte cholesterol production, and that the observed endothelial phenotype might simply be a response to changing cholesterol levels in the brain. To test whether PLX5622 affects cholesterol production in other CNS cell types, we fed adult wildtype mice control or PLX5622 diet for one month and performed RNA sequencing on acutely purified mRNA from a mixed population of astrocytes, oligodendrocytes, and neurons (**Fig 3A)**. We found no significant differences in expression of cholesterol genes between mice on control and PLX5622 diet (**Fig 3B**). These data suggest that PLX5622 increases cholesterol synthesis and uptake in endothelial cells but not in the major cholesterol-producing cells in the brain.

**Figure 3.**
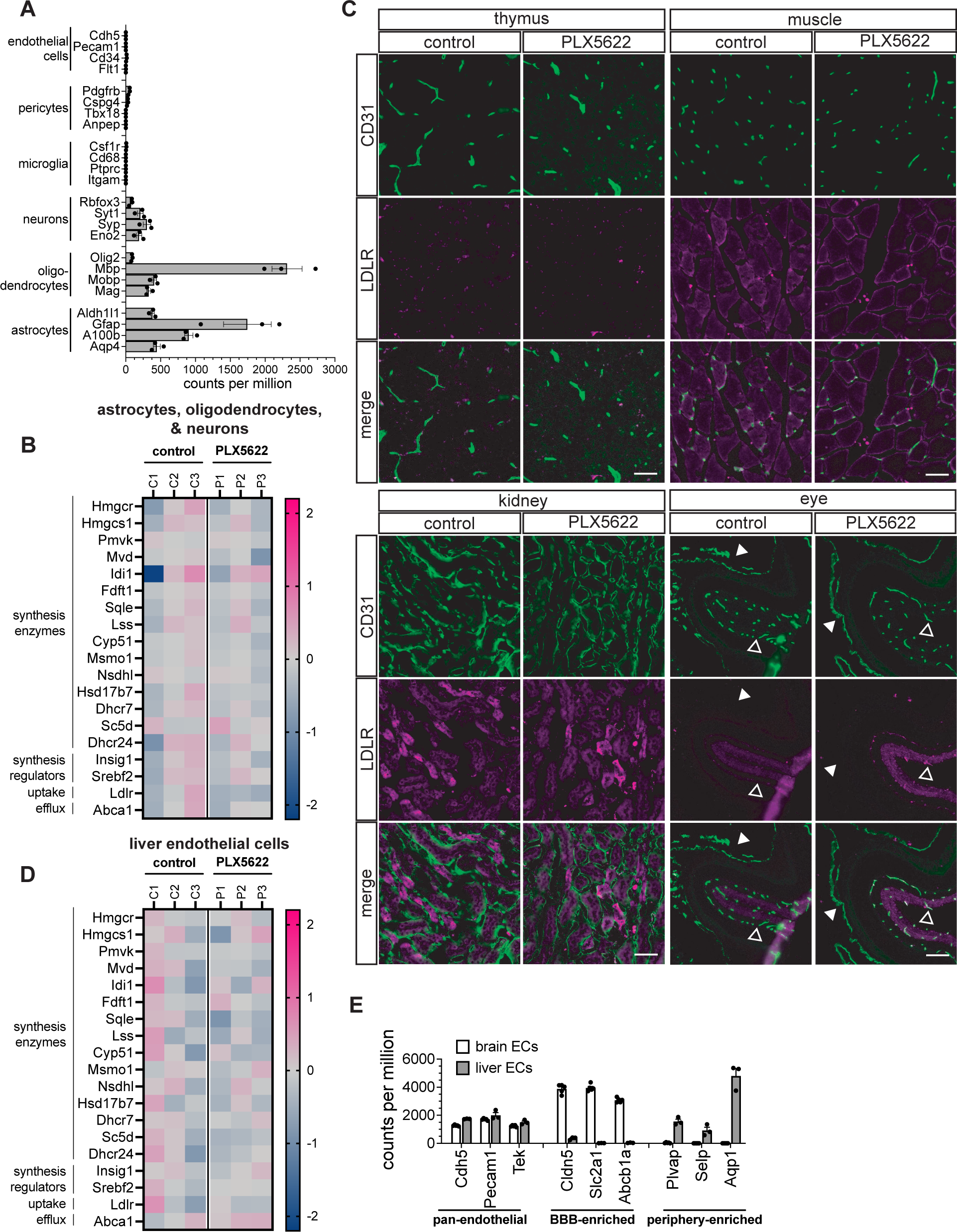
PLX5622 increases expression of cholesterol synthesis and uptake genes specifically in central nervous system endothelial cells. (A) Gene expression profile of cell-type markers in a mixed population of astrocyte, oligodendrocyte, and neuronal mRNA isolated from brains of adult mice fed control diet for one month. There was very little expression of endothelial, pericyte, or microglial markers in this neuroglial population. (B) Cholesterol-related gene expression in the mixed population of astrocytes, oligodendrocytes, and neurons from brains of adult mice fed control or PLX5622 diet for one month. Expression is represented as log_2_(FC) from the average of expression in control samples. Pink signifies increased expression and blue signifies decreased expression. PLX5622 treatment did not affect cholesterol-related gene expression in this mixed population. (C) Representative images of vascular LDLR in thymus, muscle, kidney and eye in mice fed control or PLX5622 diet for one week (thymus, muscle, kidney) or one month (eye). 10 μm-thick sections from different organs were stained with antibodies against CD31 (green) and LDLR (magenta). PLX5622 did not increase vascular LDLR in the thymus, muscle, kidney, or the non-CNS choroidal vasculature in the eye. Conversely, vascular LDLR was increased in the CNS retinal vasculature of the eye. Scale bars represent 50μm. (D) Cholesterol-related gene expression in liver endothelial cells isolated from mice fed control or PLX5622 diet for one month. Expression is represented as log_2_(FC) from the average of expression in control samples. Pink signifies increased expression and blue signifies decreased expression. PLX5622 treatment did not affect cholesterol-related gene expression in liver endothelial cells. (E) Expression of pan-endothelial, BBB-enriched, and periphery-enriched genes in the control population of liver endothelial cells shown in (D) compared to the control population of BBB endothelial cells shown in Fig 1H and 2A. As expected, BBB endothelial cells (white bars) exhibited high expression of pan-endothelial and BBB-enriched genes. Liver endothelial cells (grey) exhibited high expression of pan-endothelial and periphery-enriched genes. Expression shown in counts per million. Error bars represent SEM.

### PLX5622 alters cholesterol metabolism exclusively in CNS endothelial cells

After finding that the effect was specific to endothelial cells in the brain, we wanted to understand whether PLX5622 affects cholesterol metabolism in endothelial cells throughout the body. We fed adult wildtype mice control or PLX5622 diet for one week and examined tissue from the thymus, muscle, kidney, spinal cord, and eye. We found that PLX5622 increased vascular LDLR in the spinal cord (not shown) and retina similarly to in brain (**Fig 3C**). However, PLX5622 did not increase vascular LDLR in the thymus, muscle, kidney, or non-CNS choroidal vasculature of the eye (**Fig 3C**), suggesting that the effect is specific to CNS vasculature. To further confirm this finding and test whether it also extended to the cassette of synthesis enzymes, we fed adult wildtype mice control or PLX5622 diet for one month, acutely isolated liver endothelial cells, and performed RNA sequencing. As expected, liver endothelial cells showed high expression of known pan-endothelial and peripherally enriched genes but not BBB-enriched genes (**Fig 3E**). PLX5622 did not induce significant changes in the expression of cholesterol genes in liver endothelial cells (**Fig 3D**), demonstrating that PLX5622 modulates cholesterol-related genes specifically in CNS endothelial cells and not in endothelial cells in peripheral organs.

### Alteration in brain endothelial cholesterol synthesis is not an indirect result of neuronal hyperexcitability

Microglia have been shown to regulate neuronal activity, with PLX5622-mediated microglial depletion leading to neuronal hyperexcitability^10,11^. We wanted to address whether PLX5622 might be affecting endothelial cholesterol metabolism indirectly by increasing neuronal activity. To accomplish this, we mined a previously generated dataset in which we used chemogenetics to activate or silence neuronal activity to identify activity-dependent changes to brain endothelial cell gene expression^28^. We examined the effect of PLX5622 on the 53 endothelial genes most strongly bidirectionally regulated by neuronal activity^28^. Expression of only 3 of these activity-dependent genes was significantly altered by PLX5622, and *Sqle*, the only cholesterol synthesis gene on the list, was the only gene significantly altered by PLX5622 in the same direction as neuronal activation (**Fig 4A**). Thus, PLX5622 does not have the same broad effect on brain endothelial cells as increasing neuronal activity; it is much more specific to modulating cholesterol.

**Figure 4.**
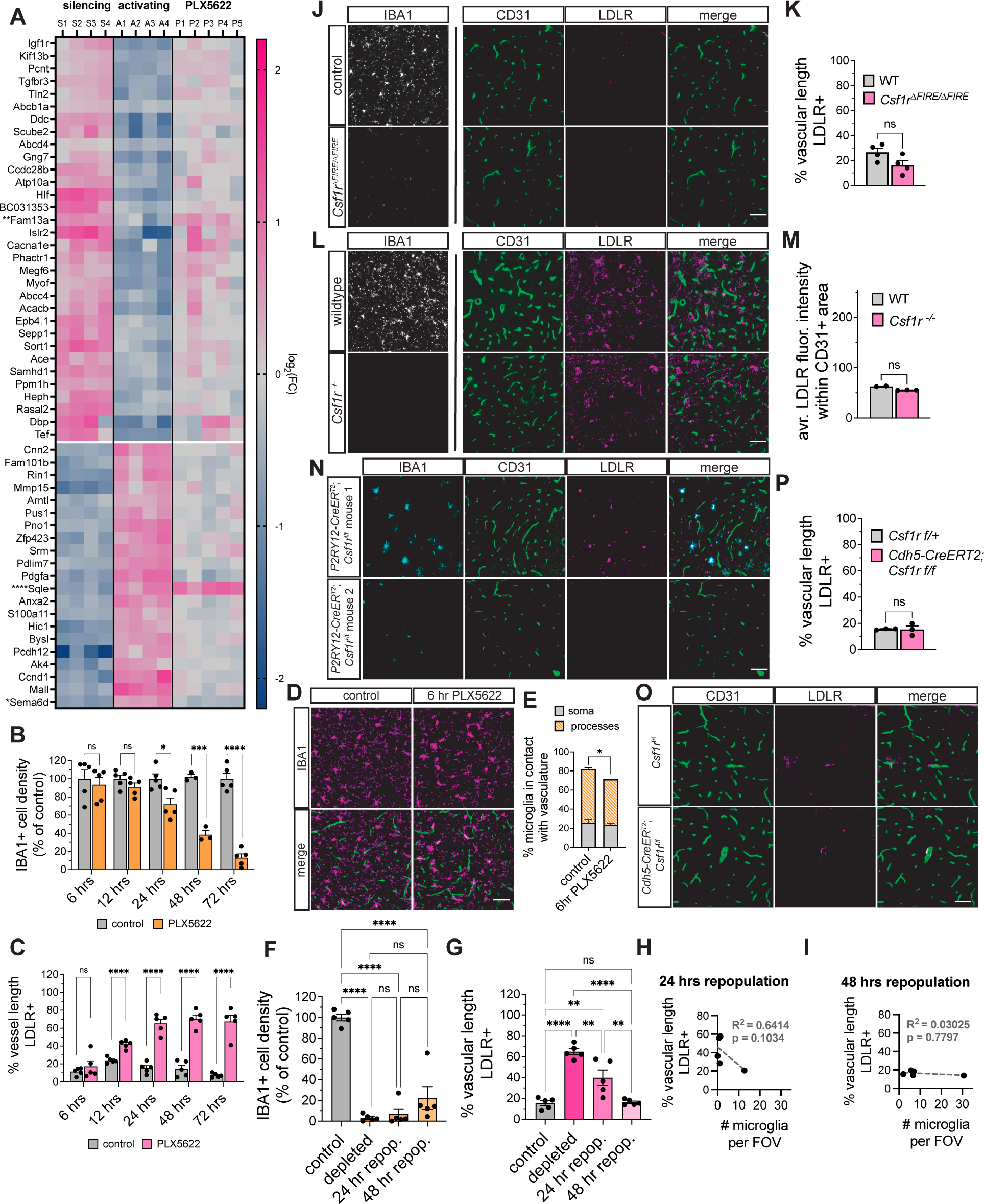
PLX5622-mediated increase in LDLR expression is independent of microglial depletion. (A) Expression of genes most strongly bidirectionally regulated by neuronal activity in activating, silencing, and PLX5622 diet conditions. Expression is presented as log2(FC) from the average of each condition’s respective controls. PLX5622 did not phenocopy neuronal activation. Only *Fam13a*, *Sqle*, and *Sema6d* were significantly altered by PLX562 (p-adj=0.00275; p-adj=4.64×10^-^15; p-adj=0.0290), and only *Sqle* was significantly altered in the expected direction. Neuronal activity data from Pulido et al. [28] (B) Quantification of microglial density in 8- to 12-week-old C57BL/6 fed control or PLX5622 diet for 6, 12, 24, 48, or 72 hours. Cortical IBA1+ cell bodies per image were counted. Each image came from a distinct 10μm section. Data are represented as a percentage of microglia normalized to the average of control. Significant microglial depletion is evident 24 hr after diet onset. n=5 mice per group per timepoint; 6 hr: p=0.622; 12 hr: p=0.181; 24 hr: p=0.0127; 48 hr: p=0.000268; 72 hr: p<0.0001, unpaired two-tailed t-tests. Error bars represent SEM. (C) Quantification of vascular LDLR in the same samples analyzed in (B). Sections were stained for LDLR and CD31. The percent CD31+ vascular length that was also LDLR+ was quantified from cortex. PLX5622 increases LDLR+ vascular length starting at 12 hr after diet onset. n=5 mice per group per timepoint, 3 sections per mouse; 6 hr: p=0.386; 12 hr: p<0.0001; 24 hr: p<0.0001; 48 hr: p<0.0001; 72 hr: p<0.0001, unpaired two-tailed t-tests. Error bars represent SEM. (D) Representative images of cortex from mice on control or PLX5622 diet for 6 hrs. Sections were stained for CD31 (green, endothelial cells) and IBA1 (magenta). Scale bar represents 50 μm. (E) Quantification of the number of microglia contacting CD31+ vasculature in the cortex at the 6 hr diet timepoint (images shown in D). The number of microglia soma and processes in contact with vasculature was divided by the total number of microglia per image to calculate the percentage of microglia making vascular contact. After 6 hr of PLX diet, significantly fewer microglia were making contact with vasculature via their processes. n=3 mice per group, p=0.0162, unpaired two-tailed t-test. Error bars represent SEM. (F) Quantification of cortical microglial density during microglial repopulation. 8- to 12-week-old C57BL/6 mice were put on control or PLX5622 diet for two weeks. Some of the PLX5622 group was then switched to control diet, while the rest remained on PLX5622 diet. Samples were taken at 24 hr and 48 hr after PLX5622 removal, creating 4 groups: control, PLX5622, 24hr repopulation, 48 hr repopulation. 10μm sections were stained with IBA1, and IBA1+ cell bodies in per cortical image were counted. Data are represented as a percentage of microglia normalized to the average of control. Microglia numbers were significantly lower in the depletion group and both repopulation timepoints compared to control, all p<0.0001. There were no significant differences between any of the three other groups. n=5 mice per group. Depleted vs 24 hr: p=0.978; depleted vs 48 hr: p=0.183; 24 vs 48 hr: p=0.336; one-way ANOVA with Tukey’s multiple comparisons test. Error bars represent SEM. (G) Quantification of vascular LDLR in the samples described in (F). 10μm sections of were stained for CD31 and LDLR. The percent CD31+ vascular length that is LDLR+ was calculated in images of cortex. LDLR levels were significantly increased from control in the depletion group (p<0.0001) and at 24 hr after PLX5622 removal (p=0.0038). Compared to the depletion group, there was significantly less LDLR+ vasculature at 24 hr (p=0.0032) and 48 hr (p<0.0001) after PLX5622 removal. By the 48 hr timepoint, the percent LDLR+ vasculature was not different from control (p=0.999). One-way ANOVA with Tukey’s multiple comparison test. Error bars represent SEM. (H) Linear regression analysis of vascular LDLR and microglial density at 24 hours from data shown in (F-G). There was no significant linear correlation between the number of microglia and the LDLR+ vascular length in cortex (n=5, p=0.103, simple linear regression). (I) Linear regression analysis of vascular LDLR and microglial density at 48 hours from data shown in (F-G). There was no significant linear correlation between the number of microglia and the LDLR+ vascular length in cortex (n=5, p=0.780, simple linear regression). (J) Representative images of cortex from 4-5 month old Csf1r^ΔFIRE/ΔFIRE^ mice and wildtype controls stained with antibodies against IBA1 (grey) or LDLR (magenta) and CD31 (green). Microglia are completely absent from Csf1r^ΔFIRE/ΔFIRE^ mice. Scale bars represent 50 μm. (K) Quantification of images shown in (J). There was no significant difference in percent CD31+ vascular length that was also LDLR+ between control and FIRE mice. n=4 mice per group, 3 sections per mouse. p=0.1143, Mann-Whitney test. Error bars represent SEM. (L) Representative images of cortex from P17-20 Csf1r^-/-^ and wildtype littermate controls stained with antibodies against IBA1 (grey) or LDLR (magenta) and CD31 (green). Microglia are completely absent from Csf1r^-/-^ mice. Scale bars represent 50 μm. (M) Quantification of images shown in (L). ImageJ was used to quantify the LDLR fluorescence intensity within thresholded CD31+ vascular area. There was no significant difference between wildtype and Csf1r^-/-^ mice. n=2-3 mice per group, 3 sections per mouse. p=0.200, Mann-Whitney test. Error bars represent SEM. (N) Representative images of cortex from two different 8-week-old P2RY12-CreER^T^^2^; Csf1r^f/f^ mice after 8 days of tamoxifen administration. Sections were stained with antibodies against IBA1 (cyan), LDLR (magenta), and CD31 (green). Level of microglia depletion varied, but vascular LDLR levels remained low. Visible LDLR instead co-localized to dying microglia. Scale bars represent 50 μm. (O) Representative images of cortex from *Cdh5-CreER^T^*^2^*; Csf1r^f/f^* and *Csf1r^f/+^* littermate controls one month after tamoxifen administration. Sections were stained with antibodies against LDLR (magenta) and CD31 (green). Scale bars represent 50 μm. (P) Quantification of images shown in (O). There was no significant difference between control and conditional knockout mice. n=3 mice per group, 3 sections per mouse. p>0.999, Mann-Whitney test. Error bars represent SEM.

### Microglial depletion does not mediate changes in endothelial cholesterol metabolism

Our first hypothesis was that microglia are required for homeostatic control of vascular cholesterol synthesis and uptake, and our finding that the cholesterol effect was specific to CNS endothelial cells fit with this idea. However, we recognized the possibility that PLX5622 might be acting independently of microglial depletion. To understand the time course of the LDLR phenotype and how it coincided with that of microglial depletion, we analyzed brain tissue sections from mice on PLX5622 diet for 6, 12, 24, 48, and 72 hours. Surprisingly, vascular LDLR was significantly increased as early as 12 hours after diet onset (**Fig 4C**). At this timepoint, there was not yet a significant reduction in microglia (**Fig 4B**), suggesting that PLX5622 acts directly on endothelial cells. However, microglial morphology is clearly affected just 6 hours after diet onset, with microglia appearing less ramified (**Fig 4D**). Indeed, after 6 hours on diet, we found that microglia made significantly fewer contacts with vasculature (p=0.0162) (**Fig 4E**). This difference was driven by the retraction of microglial processes rather than any change in somatic-vasculature contacts (**Fig 4E**). Thus, we reasoned that microglia—while not yet dead—might lose some of their homeostatic roles in the first few hours of PLX5622 treatment.

To further test whether the cholesterol phenotype was microglia-mediated, we first fully depleted microglia by treating adult wildtype mice with PLX5622 or control diet for two weeks. We then switched some of the PLX5622 group back to control diet and subsequently quantified vascular LDLR at acute timepoints of microglial repopulation. Microglia density was not significantly increased from depleted levels at either 24 or 48 hours after PLX5622 removal (**Fig 4F**). At 24 hours post PLX5622 removal, there was a large variation in vascular LDLR, with some mice still expressing high vascular LDLR levels and others already showing a large decrease (**Fig 4G**). To our surprise, by 48 hours after PLX5622 removal, levels of vascular LDLR had fully returned to control levels, despite microglia density still being at only approximately 20% of control density (**Fig 4F-G**). Among individual animals, there was no significant linear correlation between microglia number and vascular LDLR at either 24 or 48 hours after PLX5622 removal (p=0.103; p=0.780) (**Fig 4H-I**). Together, the acute depletion and repopulation experiments suggest that the effect of PLX5622 on brain endothelial cholesterol metabolism is not mediated through microglia: vascular LDLR is upregulated before microglia are eliminated and is back to control levels before microglia levels reach even a quarter of their previous density.

To further confirm that microglia do not regulate vascular cholesterol metabolism, we quantified vascular LDLR in genetic microglial depletion models. If PLX5622-mediated increase in vascular LDLR is indeed independent of microglial depletion, genetic models of microglial depletion would not exhibit changes in vascular LDLR. We first assessed vascular LDLR in *Csf1r*^ΔFIRE/ΔFIRE^ mice, which have no microglia^26^, and found no significant difference between wildtype and FIRE mice (**Fig 4J-K**). We also tested *Csf1r^-/-^* mice, which also do not possess microglia. Because these mice do not survive until adulthood, we analyzed knockouts and wildtype littermates at P20. In the developing brain, there was more extensive non-vascular LDLR in both genotypes compared to adults, but there was no increase in vascular LDLR in the knockout (**Fig 4L-M**). Finally, we looked at adult *P2ry12-CreER^T^*^2^*;Csf1r^f/f^*mice, in which tamoxifen leads to the excision of *Csf1r* from microglia, causing microglial death. The conditional knockouts had varying levels of microglial depletion, but all had low levels of vascular LDLR (**Fig 4N**). While there was bright LDLR signal in images from some mice, it was localized to dying microglia rather than endothelial cells (**Fig 4N**). These data further confirm that the observed effects of PLX5622 on endothelial cholesterol metabolism are independent of microglial depletion.

While CSF1R expression in the brain is largely limited to microglia^29,30^, some studies have suggested a functional role for endothelial CSF1R^27,31^. To determine whether the cholesterol phenotype was driven by PLX5622-mediated blockade of endothelial CSF1R, we made a tamoxifen-inducible, endothelial-specific *Csf1r* knockout mouse, *Cdh5-CreER^T^*^2^; *Csf1r*^f/f^. There was no difference in LDLR expression between knockout mice and littermate controls (p>0.999) (**Fig 4O-P**), indicating that PLX5622 does not affect brain endothelial cholesterol metabolism through inhibition of endothelial CSF1R. Taken together, these data demonstrate that PLX5622 alters brain endothelial expression of cholesterol synthesis and uptake in a manner independent from microglial depletion or endothelial CSF1R inhibition.

## Discussion

The neurovascular unit includes endothelial cells, the pericytes and smooth muscle cells that lie across the endothelial basement membrane, and the astrocytes that ensheath the vasculature with their endfeet. Diagrams of the neurovascular unit are sometimes devoid of microglia, and in other cases microglia are depicted floating in the nearby parenchymal space. However, a subset of microglia intimately associates with blood vessels in the healthy brain; either their somas contact the vasculature, or they stick one of their dynamic processes in between astrocyte endfeet to touch the endothelial basement membrane^13,15,16^. There is still much to understand about the functional relevance of these contacts.

While pericytes play a functional role in generating and maintaining BBB properties^32^, it was unknown whether microglia might influence barrier properties. Elmore et al. reported that microglial depletion does not disrupt the BBB^25^, but their methodology—dye injection and imaging of the whole brain from above—was likely not sensitive enough to detect small changes in vascular permeability. Furthermore, the BBB is not simply “open” or “closed”. There are a series of BBB properties that might be altered without increasing permeability to non-specific molecules. For instance, a change in specialized transport of particular ions or nutrients would not lead to increased extravasation of injected tracers but would still alter the extracellular environment of the CNS.

Thus, to answer the question of whether microglia play a functional role at the healthy BBB, we depleted microglia and assessed BBB ultrastructure, permeability, and expression of genes associated with BBB properties. We found no significant differences in tight junction length, number of vesicles, permeability to different types of molecules, or expression of genes associated with junctional proteins, efflux transporters, specialized nutrient transporters, or leukocyte adhesion molecules. Further, we found no differences in the expression of genes associated with BBB properties, helping to confirm our structural and functional data. It is important to note that while we found no functional role for microglia in maintaining the healthy BBB, microglia might still play a role in dynamic regulation of BBB properties in response to certain stimuli. Similarly, brain vasculature has many roles beyond its barrier properties, and our results do not preclude other functional roles for microglia-endothelial interaction. For example, recent studies suggest that microglial P2RY12 signaling is involved in neurovascular coupling^15,22^. More work is needed to fully explore microglia-vascular interactions in the healthy CNS.

Despite finding no changes to BBB properties, to our surprise, we found that PLX5622 alters brain endothelial cholesterol metabolism. PLX5622 is touted as very specific for CSF1R and has a ten-fold lower affinity for related kinases compared to PLX3397, another CSF1R inhibitor used to deplete microglia^33^. However, it is reported that the standard high dose of 1200 mg/kg PLX5622 in rodent chow leads to a concentration of 22 μM and 6.04 μM of the drug in blood and brain parenchyma, respectively^34^. These concentrations are both far above PLX5622’s reported IC_50_ values for FLT3, KIT, AURKC, and KDR^34^. Thus while PLX5622 might be more specific than PLX3397, the PLX5622 dose often used for microglial depletion is likely inhibiting other kinases and altering biological function in cell types aside from microglia. While many microglial depletion studies have validated their findings using genetic models and alternative methods to link their phenotypes to microglia, others may have erroneously connected their results to microglial depletion. Thus, the data in the present study warrant re-visiting some studies using PLX3397 or PLX5622.

## Acknowledgements

We want to thank Tara Rambaldo and Neil Sekiya at the VA Flow Cytometry CORE Research Facility and Dr. Kristen Jepsen and staff at the UCSD IGM Genomics Center for their support of this work. We would like to thank Plexxikon Inc. for supplying PLX5622 for this study. We would also like to thank Ana Williams, John Ferrer, and staff at the Animal Care Program. Samples in this manuscript were run on Illumina NovaSeq 6000 that was purchased with funding from a National Institutes of Health SIG grant (#S10 OD026929). C.P.P. was funded by an NSF GRFP and an NIH F31 (5-F31-NS-110403-2). A.L. was supported by an NIH T32 (T32-NS121727-01). A.L. was supported by an NIH T32 (T32-NS121727-01).

D.A.L. was supported by an NIH R01 (5R01CACA237376-03). F.C.B. was funded by a research grant from the Whitehall Foundation. R.D. was funded by an NIH R21 (R21-AG067035).

## Author Contributions

Conceptualization, C.P.P, R.D., and D.D.; Methodology, C.P.P., G.L.M, T.D.A, and R.D.; Software, S.S.H; Validation, C.P.P and K.B.; Formal Analysis, C.P.P, S.S.H, and K.B., Investigation, C.P.P, K.B., T.Z.Z, A.Z.Z., K.M.N, G.L.M, A.L., and T.P.M; Resources: F.C.B, T.D.A, and D.A.L; Data Curation, C.P.P and S.S.H; Writing—Original Draft, C.P.P and R.D.; Writing—Review and Editing, C.P.P. and R.D.; Visualization, C.P.P, S.S.H, and K.B; Supervision, C.P.P., R.D., F.C.B, T.D.A, and D.A.L.

## Methods

### Animals

8- to 12-week-old male C57BL/6 mice were purchased from Charles River Laboratories or Jackson Laboratories and housed at the University of California, San Diego. PLX5622 was provided by Plexxikon, Inc. or ordered from MedChemExpress and incorporated into the AIN-76A rodent diet (D10001i) at 1200 mg/kg by Research Diets, Inc. AIN-76A diet was used as a control. The diets were nutritionally identical and differed only in the presence or absence of PLX5622. Both diets were irradiated prior to use. For acute PLX5622 diet timepoints <24 hours, onset of diet occurred at 6pm at the beginning of the mice’s dark (waking) cycle. All experiments were approved by the Institutional Animal Care and Use Committee at UCSD.

For astrocyte mRNA isolation, RiboTag mice (*Rpl22*^tm1.1Psam^/SjJ)––which contain a ribosomal protein (Rpl22) with a floxed C-terminal exon followed by an identical exon tagged with hemagglutinin (HA)—were mated to mice in which Cre-recombinase is driven by GFAP (B6.Cg-Tg(Gfap-cre)73.12Mvs/J). Both mouse lines were purchased from Jackson Labs (#029977; #012886). 8-week-old mice were put on control or PLX5622 diet for one month before RNA isolation. All experiments were approved by the Institutional Animal Care and Use Committee at UCSD.

*Csf1r*^ΔFIRE/ΔFIRE^ mice^26^ were housed at University of California, Irvine. Mice were 4-5 months old at time of sample collection. Control samples for *Csf1r*^ΔFIRE/ΔFIRE^ mice were age-matched wildtype mice from the same line. Experiments were approved by the Institutional Animal Care and Use Committee at UCI.

*P2ry12-CreER^T^*^2^*; Csf1rf^/f^*mice were housed at the University of California, San Francisco. 8-week-old mice were given eight days of tamoxifen treatment (150uL of 20mg/mL in corn oil, gavage) to deplete microglia prior to tissue collection on the ninth day. Experiments were approved by the Institutional Animal Care and Use Committee at UCSF.

*Csf1r*^-/-^ mice were housed at the University of Pennsylvania. Mice were P17-20 at time of sample collection. Control samples for *Csf1r*^-/-^ mice were wildtype littermates of knockouts. Experiments were approved by the Institutional Animal Care and Use Committee at UPenn.

### Immunohistochemistry

Mice were transcardially perfused with DPBS followed by 4% paraformaldehyde in 1x PBS. Tissue was cryopreserved in 30% sucrose, frozen in 2:1 OCT:30% sucrose and sectioned at 10 μm thickness. Sections for LDLR or tight junction protein staining were further fixed in ice-cold methanol for 10 minutes. Sections for IBA1 staining were further fixed with 5 minutes of 4% PFA. All slides were blocked with 10% donkey serum, 0.2% triton-x-100 in PBS. Primary antibodies against LDLR, IBA1 (Wako), PECAM1, CLDN5, OCLN, and P-GP were used at 1:1000. Primary antibody against IBA1 (Novus Bio) was used at 1:500. Primary antibody solution was 10% blocking solution in primary antibody buffer (150 mM NaCl, 50mM Tris base, 1% BSA, 100mM L-lysine, 0.02% sodium azide in dH_2_O). Primary antibody incubation occurred overnight at 4°C. Fluorescently conjugated secondary antibodies (Life Technologies) were used at 1:1000 and incubated for 1.5 hours at room temperature. Slides were coverslipped using DAPI-Fluoromount-G (SouthernBiothech 0100-20) and imaged with a Zeiss Axio Imager.D2.

### Transmission electron microscopy (TEM)

Mice were transcardially perfused with DPBS followed by 15 mL of 2% paraformaldehyde and 2.5% glutaraldehyde in 0.15M sodium cacodylate buffer. Brains were dissected and post-fixed in the same solution overnight at 4°C. Small cubes were cut from cortex and processed by the CMM Electron Microscopy Facility. Ultrathin sections were cut on a Leica microtome with a diamond knife and stained with uranyl acetate and lead. Images were captured on a JEOL 1400plus TEM at 80 kV with a Gatan 4kx4k camera. At least 20 individual vessel cross-sections were imaged per animal.

### Tight junction and transcytosis analysis

The length of tight junctions across all TEM images was measured using the ImageJ line tool and converted to nanometers using the image scale bar. Sum length of all tight junctions was divided by the number of tight junctions to determine average tight junction length. Invaginating vesicles were counted across all images for each mouse. The total number of invaginating vesicles was divided by the number of vessel cross-sections analyzed.

### Sodium fluorescein

Mice were injected (i.p.) with 25 mg/kg sodium fluorescein salt (Sigma F6377), 2 mg/mL in PBS. After 90 minutes, blood samples were taken from the right ventricle and transferred to an EDTA-coated tube on a rotator. Mice were perfused with DPBS. Brains were dissected, meninges were removed, and cortex and hippocampus were flash frozen. Plasma was collected from blood samples and flash frozen. Samples were stored at −80°C until further processing. Tissue was homogenized in PBS, plasma was diluted in PBS, and 2% TCA was added to each to extract sodium fluorescein overnight. Samples were spun, and supernatant was collected and diluted with borate buffer. A Tecan Infinite 200 plate reader was used to measure fluorescence (excitation 480, emission 538). Wells of 0.5% TCA, 50% borate buffer, and 25% PBS were used as a background control. Average of background signal was subtracted from that of each experimental well. Brain fluorescence was divided by blood fluorescence within each sample to determine the amount of parenchymal extravasation.

### Rhodamine 123

Mice were injected with 2 mg/mL rhodamine 123 (1:4 DMSO:sterile saline) at a dose of 25 mg/kg. Mice were perfused with DPBS. Brains were dissected, meninges were removed, and cortex and hippocampus together were flash frozen. Plasma was collected from blood samples and flash frozen. Samples were stored at −80°C until processing. Tissue was homogenized in PBS, plasma was diluted in PBS. Rhodamine123 was extracted by vortexing and incubating overnight with butanol. A Tecan Infinite 200 plate reader was used to measure fluorescence (excitation 505, emission 560). Wells of butanol were used as a background control. Average of background signal was subtracted from that of each experimental well. Brain fluorescence was divided by blood fluorescence within each sample to determine the amount of parenchymal extravasation.

### Western blot

Mice on control or PLX5622 diet for one month were euthanized with carbon dioxide. Cortices were dissected and flash-frozen. Samples were thawed in RIPA buffer with protease inhibitors, and homogenized in a 1.5 mL glass dounce. Supernatant was analyzed for protein concentration using the Pierce BCA Protein Assay Kit. Samples were mixed with 6x laemmli buffer and incubated at 100°C for 7 minutes. A PageRule ladder and 15 ug of each sample was loaded on a Bolt 4-12% Bis-Tris Plus gel and run at 65V for 15 minutes, followed by 100V for 1 hr 20 min. Proteins were transferred to a nitrocellulose membrane at 120 V for 1.5 hours. Blots were blocked with 5% milk in TBST. Primary antibody incubation was performed at 4°C overnight. Antibodies against CLDN5, OCLN, and P-GP were diluted 1:1000 in 5% BSA + 0.02% sodium azide. Anti-beta-actin was diluted 1:10,000. Secondary incubation was performed at room temperature for 2 hours. Anti-mouse and anti-rabbit HRP-conjugated antibodies were diluted 1:4000 in 5% milk. Membranes were imaged using an Azure 280 imaging system (Azure Biosystems). ImageJ was used to quantify band intensity.

### qPCR

Adult mice fed control or PLX5622 diet for one month were euthanized with carbon dioxide. Brains were dissected and flash frozen. Three frozen brains per sample were thawed in a 1x HBSS buffer with HEPES and sodium bicarbonate. Tissue was minced with a #10 scalpel blade and homogenized in a glass dounce. After centrifugation, pellet was vigorously shaken with HBSS buffer containing 3.6 g/mL dextran. Samples were transferred to ultracentrifuge tubes and spun at 4400g for 15 minutes at 4C with no brake.

Myelin layer and supernatant were aspirated, and pellet was resuspended in HBSS buffer and filtered through glass beads in a 100μm filter. Vessel segments attached to the glass beads were shaken in HBSS buffer with 1% BSA to detatch vessels. Vessels were then filtered through a 100 μm filter held above a 20 μm mesh filter. The vessel segments captured by the 20 μm mesh were washed off and spun down. Pellet was rinsed with DBPS, liquid was removed, and pellets were flash frozen on dry ice. Pelleted cells were lysed with trizol and RNA isolation was performed with the Qiagen RNeasy Micro kit (74004). PrimeTime^TM^ qPCR primers for *Ldlr* (Mm.PT.58.23359070), *Dchr24* (Mm.PT.58.5182147), and *Gapdh* (Mm.PT.39a.1) were purchased from Integrated DNA Technologies. cDNA was made from vessel fraction RNA using iScript Reverse Transcription Supermix. qPCR was performed using SYBR Green Master Mix. The ΔΔCt method normalizing to *Gapdh* was used to calculate relative gene expression of *Ldlr* and *Dhcr24.* In each case, gene expression was further normalized to the average of expression in the control samples.

### Endothelial enrichment for RNA sequencing

Mice were anesthetized with a cocktail of ketamine (4 mg/ml) and xylazine (0.6 mg/ml) in saline. Mice were then decapitated, and brains or livers were dissected. For brain samples, cerebellum and olfactory bulbs were removed and discarded. Brains were rolled on filter paper to remove meninges. Remaining meninges and choroid plexus were removed with fine forceps. Brain or liver tissue was diced using a #10 scalpel blade. Tissue was enzymatically digested with the Papain Dissociation System kit (Worthington, LK003176), using 1 vial (>100 units) per 10 mL EBSS (Sigma, E7510) solution also containing 0.36% D(+)-Glucose, 26 mM NaHCO3, 0.5 mM EDTA, and 62.5 units/mL DNase (Worthington, LS002007) at 35°C for 90 minutes with 95% O_2_, 5% CO_2_ continuously passed over the solution. Tissue chunks were washed with a “low-ovomucoid” EBSS solution containing 225 μg/mL ovomucoid (Worthington, LS003089), 225 μg/mL BSA (Sigma A8806), 0.36% D(+)-Glucose, 26 mM NaHCO3, and 55.5 units/mL DNase. Samples were triturated with 10 mL, 5 mL, and P1000 pipette tips, successively. Cells were spun into “high-ovomucoid” solution containing 450 μg/mL ovomucoid, 450 μg/mL BSA, 0.36% D(+)-Glucose, 26 mM NaHCO3, and 62.5 units/mL DNase. Cells were resuspended in a collagenase/dispase solution of 1 mg/mL collagenase type II (Worthington, LS004176) and 0.4 mg/mL neutral protease (Worthington LS02104) in an EBSS solution containing 0.36% D(+)-Glucose, 26 mM NaHCO3, 0.5 mM EDTA, and 62.5 units/mL DNase. Cells were incubated for 30 min at 35°C with 95% O_2_, 5% CO_2_ continuously passed over the solution. After incubation, cells were spun into high-ovomucoid solution. Brain cells were resuspended in a 0.5% BSA (Sigma, A4161) solution with myelin removal beads (Miltenyi Biotec, 130-096-433). After a 15 min bead incubation, 30 μm pre-separation columns (Miltenyi Biotec, 130-041-407) and LS columns (Miltenyi Biotec, 130-042-401) were used for myelin removal from brain samples. Conversely, liver samples were filtered through a 30 μm filter and red blood cells were lysed with ACK buffer for 5 minutes. After myelin removal or red blood cell lysis, cells were blocked with rat IgG in 0.5% BSA solution on ice for 20 min. Cells were stained with AF647-conjugated anti-CD31 (Molecular Probes A14716), FITC-conjugated anti-CD13 (BD Biosciences 558744), FITC-conjugated anti-CD45 (eBioscience 11-0451-85), FITC-conjugated anti-CD11b (eBioscience), and DAPI. Cells positive for 647 and negative for FITC and DAPI were sorted into trizol at the UCSD Flow Cytometry Research Core Facility. RNA isolation was performed with the Qiagen RNeasy Micro kit (74004).

### RNA isolation of glial population

Adult GFAP-Cre RiboTag mice fed control or PLX5622 diet for one month were euthanized with CO_2_. A half forebrain from each mouse was dissected and homogenized in RNase-free water with 1% NP-40, 0.1M KCl, 50 mM Tris, and 12 mM MgCl_2_ supplemented with 0.1 mg/mL cycloheximide, protease inhibitors, 1 mg/mL heparin, RNAsin, and 1mM DTT. Homogenized tissue was spun down and supernatant was incubated with anti-HA (CST 3724) at 1:160 for four hours at 4°C. Magnetic IgG beads (Pierce 88847) were added and incubated overnight. Beads were washed 3 times with high salt buffer (RNase-free water with 0.3 M KCl, 1% NP-40, 50 mM Tris, 12 mM MgCl_2_, 0.1 mg/mL cycloheximide, and 0.5 mM DTT) using a magnetic stand. RLT lysis buffer + 1% βME was added to beads and vortexed. Supernatant was collected and mRNA was purified by Qiagen RNAeasy Micro kit (74004).

### Bulk sequencing and bioinformatics

RNA isolation was performed with the Qiagen RNeasy Micro kit (74004). RNA samples were processed by the UCSD Institute of Genomic Medicine Core. A tape station bioanalyzer was used to determine quality and concentration. Non-stranded cDNA libraries were made using the TruSeq RNA Library Prep Kit v2. 100 bp paired-end sequencing was performed using an Illumina NovaSeq 6000. HISAT2 v2.1.0 was used for alignment to the GRCm38 genome. htseq-count v0.9.1 was used to generate count tables. Differential expression analyses including p-value, FDR, and log2(FC) were performed using DESeq2 v1.22.1 and Excel. All analysis programs were run through the Galaxy open-source platform.

### Single-cell sequencing and bioinformatics

Gene counts were obtained by aligning reads to the mm10 genome (refdata-gex-mm10-2020-A) using CellRanger software (v.6.0.1 – 10X Genomics). Using Seurat (v.4.0), quality control was performed using the following: 1) outliers with a high ratio of mitochondrial RNAs (>5%, <200 features) relative to endogenous RNAs and homotypic doublets (>5000 features) were removed in Seurat; 2) after scTransform normalization and integration, doublets and multiplets were removed using DoubletFinder; 3) cells were manually inspected using known cell-type specific marker genes, cells expressing more than one cell-type specific marker were removed.

In Seurat (v.4.0), gene counts were first normalized using scTransform, then Integration function was used to align data with default settings. Genes were projected into principal component (PC) space using the principal component analysis function RunPCA. The first 30 dimensions were used as inputs into Seurat’s FindNeighbors and FindClusters functions. Then, RunUMAP function with default settings was used to calculate 2-dimensional UMAP coordinates and search for distinct cell populations. Distinct cell types were determined using known cell-type specific markers. Differential gene expression of genes comparing PLX to control samples was performed with the FindMarkers function, specifically utilizing the Wilcoxon Rank Sum test.

### Analysis of LDLR+ vascular length

To quantify LDLR+ vascular length, sections were stained for LDLR and CD31. LDLR signal was traced using the ImageJ line tool, with each segment saved as an ROI. ROIs were then opened on the CD31 channel, and any ROI not corresponding to CD31+ vasculature was deleted. The sum of lengths of remaining ROIs was used as the length of LDLR+ vasculature. The rest of the LDLR-/CD31+ vascular segments were then traced in the same way, and the ROIs measured. The sum total length of all segments was used as the vascular length. Percent LDLR+ vascular length was calculated as (LDLR+ vessel length)/(total vessel length)*100.

### Analysis of LDLR intensity across vascular area

To quantify LDLR intensity across vascular area, sections were stained with antibodies against LDLR and CD31. Using ImageJ, CD31 signal was thresholded, and an ROI for total vascular area was created. This ROI was then opened on the LDLR channel image, and mean fluorescence intensity was calculated.

### Microglial repopulation

For acute repopulation experiments, mice were split into three groups: control, depleted, and repopulation. Mice were kept on AIN-76A (control group) or PLX5622 (depleted and repopulation groups) diet for two weeks. After two weeks, at 6pm, mice in the repopulation group were switched to AIN-76A diet, and tissue was collected 24 or 48 hours after diet switch, along with tissue from control and depleted groups.

### Resource Availability

All sequencing data will be uploaded to Gene Expression Omnibus (GEO)

